# Training deep learning models for cell image segmentation with sparse annotations

**DOI:** 10.1101/2023.06.13.544786

**Authors:** Ko Sugawara

**Affiliations:** Institut de Génomique Fonctionnelle de Lyon (IGFL), École Normale Supérieure de Lyon, France; Centre National de la Recherche Scientifique (CNRS), France; Laboratory for Developmental Dynamics, RIKEN Center for Biosystems Dynamics Research (BDR), Kobe, Japan

**Keywords:** deep learning, segmentation, bioimage analysis, sparse annotation

## Abstract

Deep learning is becoming more prominent in cell image analysis. However, collecting the annotated data required to train efficient deep-learning models remains a major obstacle. I demonstrate that functional performance can be achieved even with sparsely annotated data. Furthermore, I show that the selection of sparse cell annotations significantly impacts performance. I modified Cellpose and StarDist to enable training with sparsely annotated data and evaluated them in conjunction with ELEPHANT, a cell tracking algorithm that internally uses U-Net based cell segmentation. These results illustrate that sparse annotation is a generally effective strategy in deep learning-based cell image segmentation. Finally, I demonstrate that with the help of the Segment Anything Model (SAM), it is feasible to build an effective deep learning model of cell image segmentation from scratch just in a few minutes.

## Introduction

Deep learning has become a powerful approach to bioimage analysis. In particular, cell image segmentation is a topic that has attracted the attention of many researchers because it is a fundamental step in many analysis workflows. StarDist (1, 2) is a widely used algorithm that was originally developed for 2D cellular images (1) and later extended to 3D images (2). Cellpose (3, 4) has recently been developed as a universally applicable cell segmentation algorithm. In these algorithms, adequate performance can be achieved by applying a versatile pre-trained model if the target image is a typical cellular image. However, it requires training a model with additional annotated data with characteristics similar to the target image when the pre-trained model is not applicable. Because the current implementation of the algorithms requires annotation to be performed on the entire region of the input images, the user has to spend considerable time on the annotation process.

On the other hand, ELEPHANT (5), an algorithm for cell tracking, is a method that can train deep learning models with sparse annotations (Fig.1). ELEPHANT performs the tracking task in two stages: cell detection and linking, where cell detection is performed internally using U-Net (6, 7) based segmentation. In ELEPHANT, training of the segmentation model is performed using labels generated from sparse ellipse (2D) or ellipsoid (3D) annotations that represents cell instances. Because the deep learning model in ELEPHANT is designed to work with sparse annotation, users do not need to annotate all cells in input images. Instead, they interactively iterate a cycle of annotation, training, prediction and proofreading. I introduced sparse annotation to StarDist and Cellpose by making their loss functions compatible with sparsely annotated labels.

**Fig. 1.**
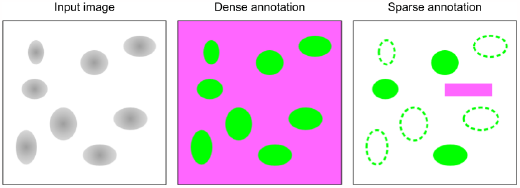
Dense annotation and sparse annotation. Foreground annotation is shown in green and background annotation in magenta.

In this work, I describe how sparse annotation-based training works with these algorithms. For each algorithm, I investigated the relationship between the number of annotations and performance and how the behavior differs with and without background pixel annotations, in addition to cell annotations. I further showed that the selection of sparse annotations can change performance, even when training is performed with the same number of annotations.

As a proof-of-concept for this project, I generate sparse annotations by running the Segment Anything Model (SAM) (8) on QuPath (9) and train the StarDist model from scratch, demonstrating that it is feasible to build an effective deep learning model of cell segmentation in just a few minutes. These sparse annotation-supported algorithms are available as open source.

## Results

### Training of deep learning models for cell segmentation with sparse annotations

A dataset with sparse cell annotations, containing 447 training data and 50 validation data, was created by the procedure described in Methods (Fig.2). Briefly, there are three annotation approaches: (i) dim nuclei preferentially, (ii) bright nuclei preferentially, and (iii) both mixed. I also included conditions with additional background labels for each of them. The number of annotations was incrementally increased to train and evaluate the deep-learning models (Fig.2).

**Fig. 2.**
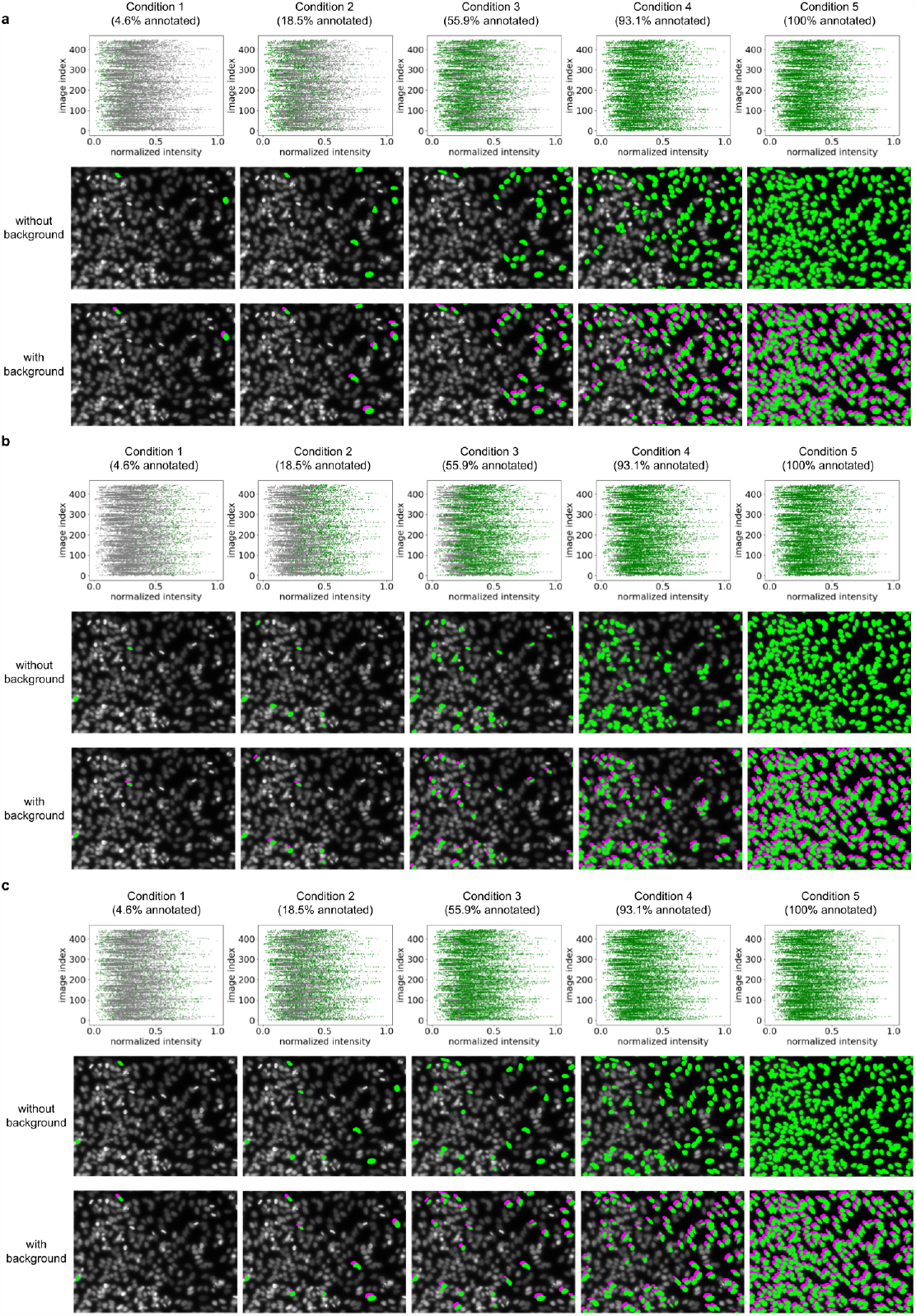
Dataset with sparse nuclei annotations. (a) Sparse annotations selected dim nuclei preferentially. (b) Sparse annotations selected bright nuclei preferentially. (c) Sparse annotations selected both mixed. The top panels of each approach show the distribution of normalized intensity values, with the labeled data points in green and unlabeled in magenta. The middle panels of each approach show the annotations overlaid on the original image, with the foreground in green. The bottom panels of each approach show the annotations overlaid on the original image, with the foreground in green and the background in magenta.

StarDist, Cellpose, and ELEPHANT models were trained using the training data under these conditions, and their performance was evaluated using the validation data. In StarDist, cell nuclei were well identified with sparse annotations, but many false positives were observed in conditions that did not include background annotations (Fig.3, 6). When dim nuclei were preferentially annotated, the Recall score increased, but the Precision score decreased. On the other hand, when bright nuclei were selected preferentially, the Recall score was low when the number of annotations was around 4.6%. When the number of annotations reached about 18.5%, the Recall score improved while maintaining the Precision score. A mixture of dim and bright nuclei showed intermediate behavior.

**Fig. 3.**
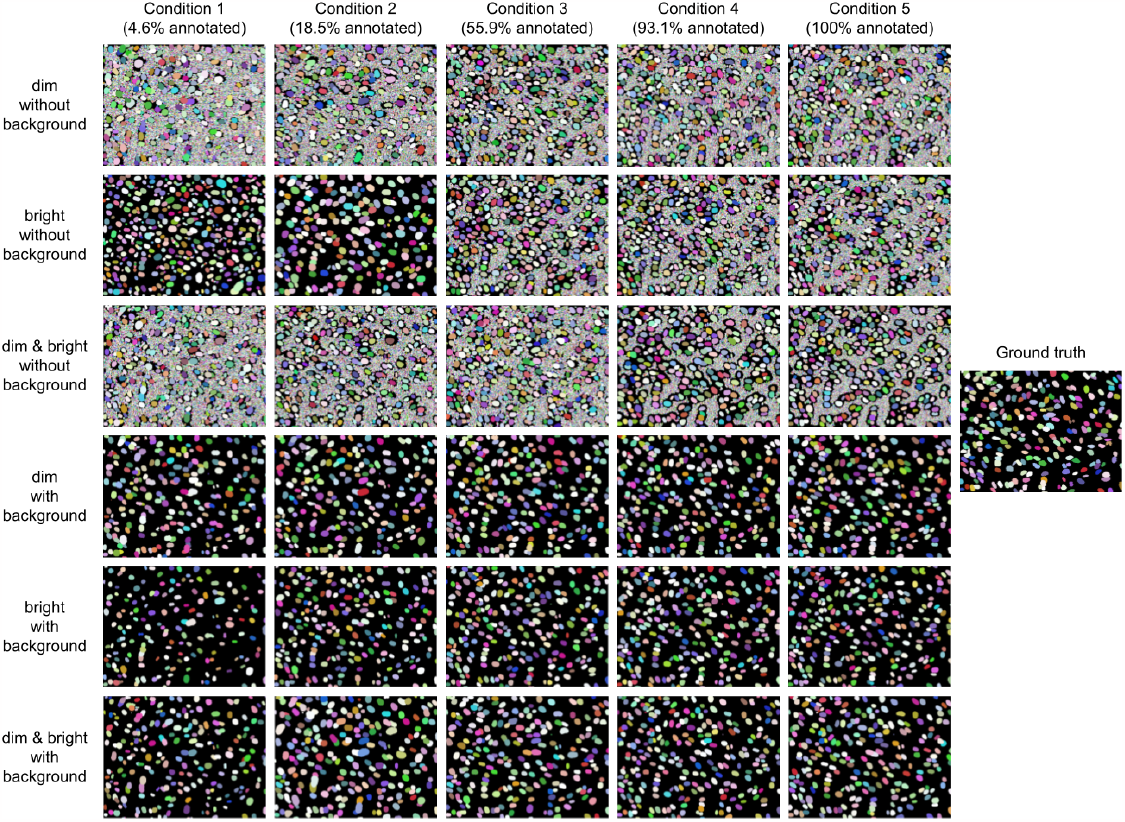
Examples of inference results of StarDist models trained under the conditions shown in each matrix. Each nucleus is colored with a randomly assigned color.

In Cellpose, the boundary between nuclei and background regions could not be discriminated in the conditions without background annotation, and both Recall and Precision showed low scores (Fig.4, 6). On the other hand, when background annotations were included, training with sparse annotations performed well. As shown in StarDist, prioritizing dim nuclei increases the Recall score, while prioritizing bright nuclei increases the Precision score, and including dim and bright nuclei exhibited intermediate properties.

**Fig. 4.**
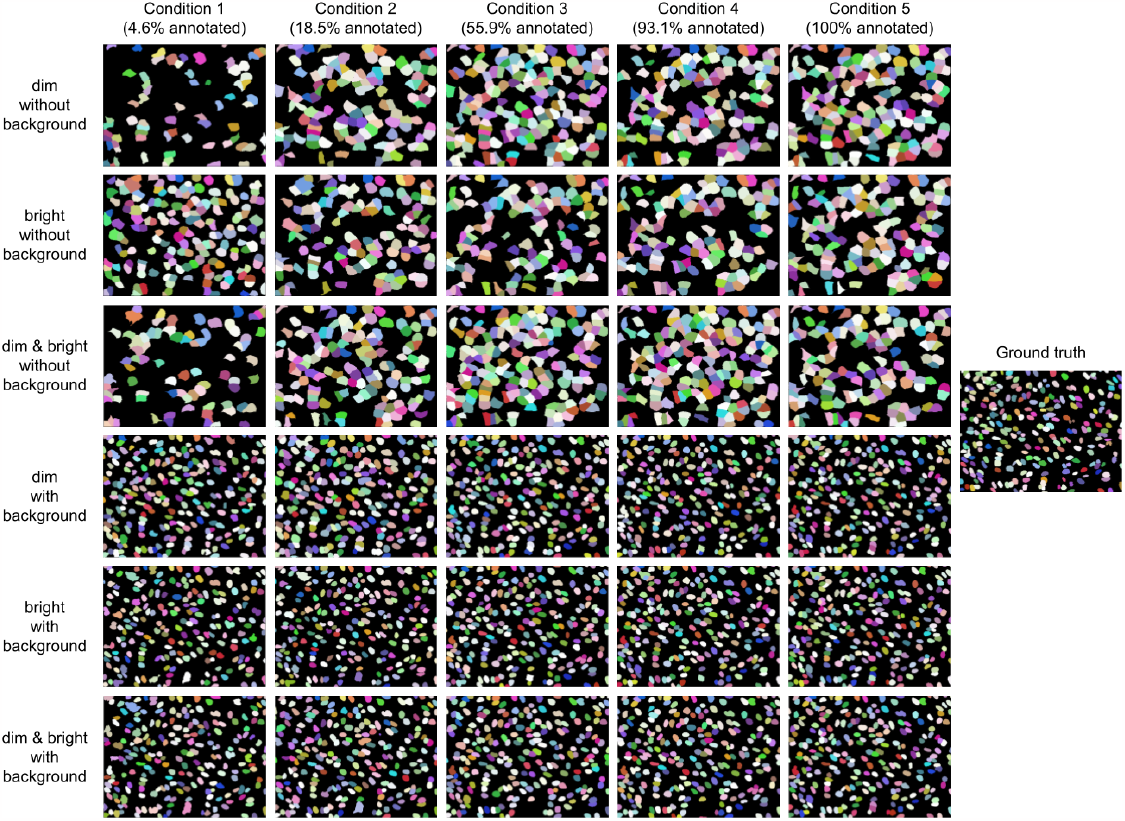
Examples of inference results of Cellpose models trained under the conditions shown in each matrix. Each nucleus is colored with a randomly assigned color.

In ELEPHANT, effective models could be built with sparse annotations, even without background annotation (Fig.5, 6). This could be due to the fact that in ELEPHANT, the foreground annotation is converted to two-layer labels: a center region and a periphery region (5). The tendency of performance with intensity-based nucleus selection was similar to StarDist, and Cellpose. Including both dim and bright nuclei seemed the best approach, especially when the number of annotations was limited.

**Fig. 5.**
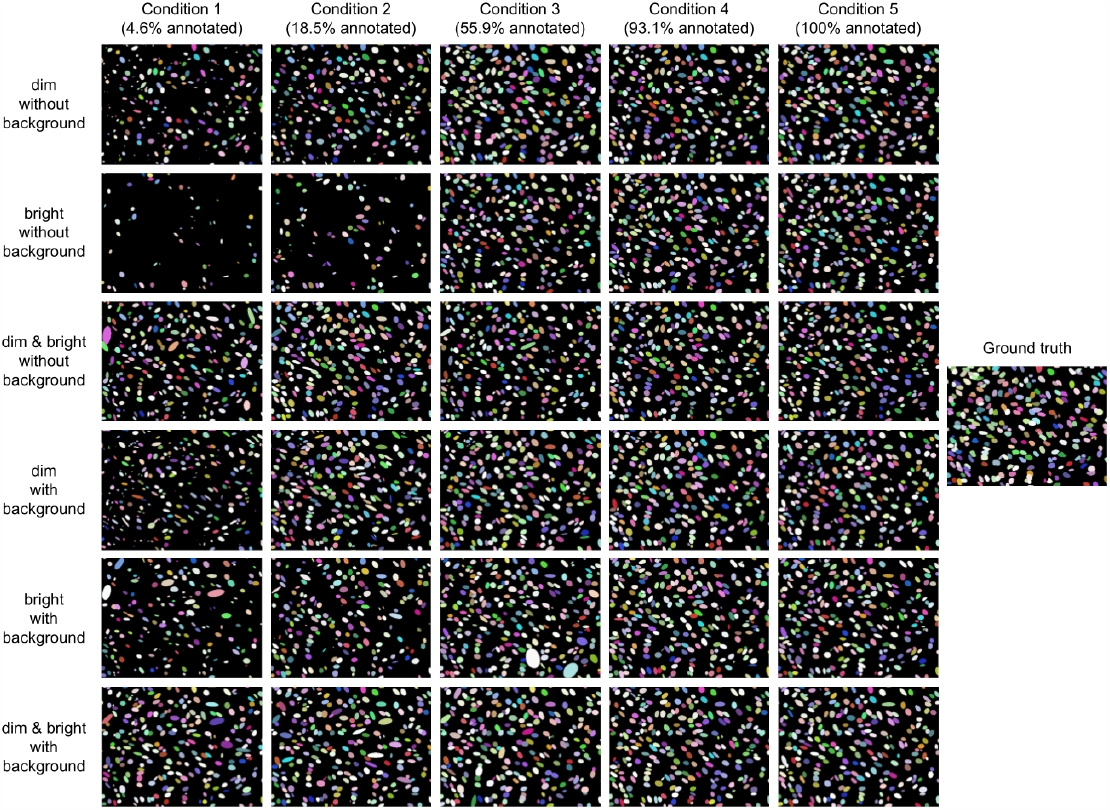
Examples of inference results of ELEPHANT models trained under the conditions shown in each matrix. Each nucleus is colored with a randomly assigned color.

**Fig. 6.**
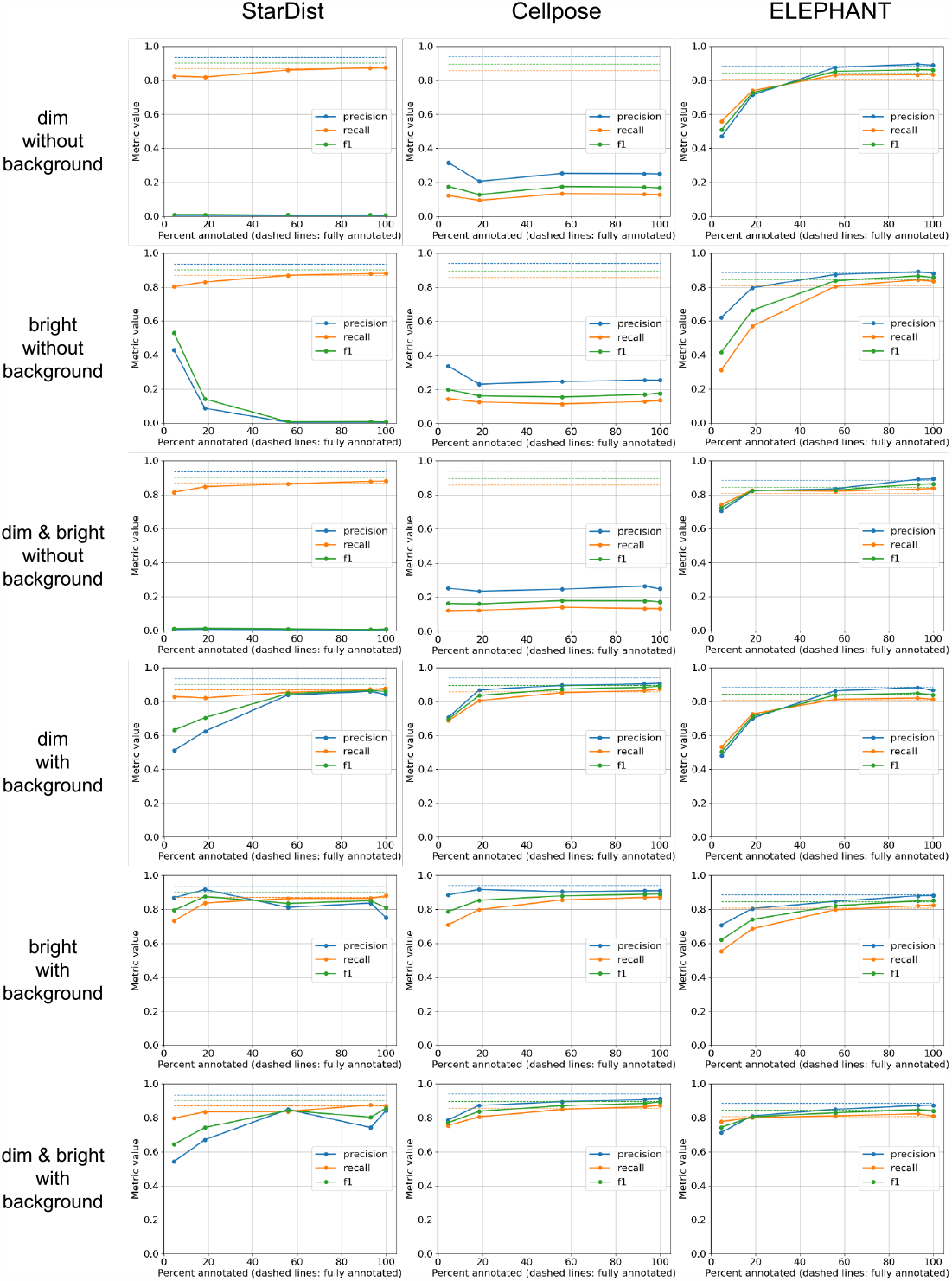
Quantitative evaluation of Recall, Precision, and F1 scores of StarDist, Cellpose, and ELEPHANT models trained under the conditions shown in each matrix. The dashed lines show the results obtained from training based on dense annotations.

### Training of a deep learning model from scratch in a few minutes

To demonstrate that training deep learning models based on sparse annotation is practical, I implemented extensions on the widely used image analysis tool QuPath (9). To facilitate the annotation process, I have implemented an extension to perform SAM (8), which enables the generation of a cell nucleus annotation from a bounding box. In addition, I implemented an extension that calls sparse annotation-based training and inference with a StarDist model. The training was performed from scratch with no prior training. I confirmed that an effective StarDist model can be trained from a few sparse annotations created by SAM in a few minutes. (Supplementary video 1: https://doi.org/10.5281/zenodo.8020156).

## Discussion

This study showed that background annotation is essential for training the StarDist and Cellpose models. Introducing the embedding consistency loss used in (10) may enable training them without background annotation. However, a major concern is that it requires another convolutional neural network in the training phase, which would increase the computational cost.

## Conclusion

In this study, I explored sparse annotation-based training of deep learning models for cell image segmentation. Widely used cell segmentation algorithms, StarDist and Cellpose, were made compatible with sparse annotation. I trained the models of the modified versions of StarDist and Cellpose, and ELEPHANT’s segmentation module with sparse annotations generated under multiple conditions, including three strategies: (i) dim nuclei preferentially, (ii) bright nuclei preferentially, and (iii) both mixed, and five annotation percentages ranging from 4.6% to 100%. I showed that for training StarDist and Cellpose models, background information is essential. I found that recall and precision scores changed depending on the annotation strategy; prioritizing dim nuclei produced higher recall scores, and prioritizing bright nuclei produced higher precision scores. The results showed models trained with relatively small sparse annotations (4.6% to 18.5%) performed similarly to fully annotated data. Finally, I demonstrated that with the help of SAM, an efficient StarDist model was trained from scratch with a few sparse annotations just in a few minutes.

## Methods

### Adaptation of StarDist and Cellpose algorithms to sparse annotation

tarDist and Cellpose are widely used algorithms, but require densely annotated data for training with custom data. I adapted these algorithms to sparse annotation with the same idea as implemented in ELEPHANT. The modification is very simple: the loss function is calculated by ignoring unannotated pixels in the training phase. These changes can be found in the following GitHub repositories (https://github.com/ksugar/stardist-sparse and https://github.com/ksugar/cellpose-sparse).

### Dataset preparation

I used a dataset from the StarDist paper (1). The dataset contains 447 training data and 50 validation data, which are a subset of the Data Science Bowl 2018, an annotated collection of 2D nuclear images acquired by fluorescence microscopy. In addition to the original fully annotated data, sparsely annotated data were prepared for the evaluation in this study. In generating the sparsely annotated data, a maximum of 2n nuclei (*n* = 1, 4, 16, 64, 256) were selected in each image, not exceeding the number of nuclei in the image, under the following three conditions: (i) 2n nuclei with the lowest intensities *(min)*, (ii) 2n nuclei with the highest intensities *(max)*, and (iii) n nuclei with the lowest intensities and n nuclei with the highest intensities *(min-max)*. Cell instances with a minor axis calculated as zero were excluded. Here, the cell annotations were prioritised over the background annotations if they are overlapped, and annotated regions outside the image area were ignored.

### Evaluation of performances

Performance evaluation was carried out on the validation data using the evaluation method in the StarDist library (1). In this method, the detected cell instances are matched with the groud-truth cell instances and if their intersection over union (IoU) exceeds a threshold value, they are counted as *true positive (TP)*, otherwise they are counted as *false positive (FP)*. After the matching, the remaining ground-truth instances were counted as *false negative (FN)*. In this study, the threshold was set at 0.5 and 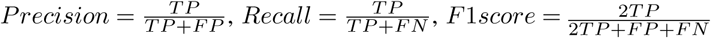 were adopted as the evaluation metric.

### Training of deep learning models

The following data augmentations were stochastically applied to input images during training: rotation, resize, horizontal/vertical flip, intensity modification. The batch size was set to 8, the number of steps per epoch to 56, training was performed for 100 epochs, and the model from the last epoch was adopted.

### Computer setup and specifications

In this study, I used a desktop computers (Dell Precision 7920 Tower Workstation or Dell Precision 3660 Tower Workstation) with the following specifications: 2x Intel Xeon Gold 5220R CPU 2.2GHz, 12×32 GB DDR4 2933 MHz RAM, Dual NVLink Nvidia RTX A6000 48GB 4DP, M.2 2TB PCIe NVMe Class 40 Solid State Drive and 14 TB external Hard Disk Drive, Ubuntu 20.04 (Dell Precision 7920), and Intel Core i9-12900 CPU 2.4GHz, 4×32 GB DDR5 3600 MHz RAM, Nvidia RTX A4000 16 GB, M.2 1TB PCIe NVMe Class 40 Solid State Drive and 4 TB Hard Disk Drive, Windows 11 (Dell Precision 3660).

## Data availability

The subset of DSB2018 dataset used in this study is from the StarDist paper (1), available at https://github.com/stardist/stardist/releases/download/0.1.0/dsb2018.zip.

## Code availability

The implementation of StarDist with support for sparse annotation is available at https://github.com/ksugar/stardist-sparse. The implementation of Cellpose with support for sparse annotation is available at https://github.com/ksugar/cellpose-sparse.

Programs for training and evaluation of StarDist, Cellpose, and ELEPHANT with sparse annotations are available at https://github.com/ksugar/cellsparse-core, https://github.com/ksugar/cellsparse-api and https://github.com/ksugar/qupath-extension-cellsparse.

The implementation of SAM for QuPath is available samapi at and https://github.com/ksugar/samapi and https://github.com/ksugar/qupath-extension-sam.

## Acknowledgements

I am grateful to Michalis Averof (IGFL, CNRS) in whose lab this work was initiated and carried out, and to Shuichi Onami (RIKEN BDR) in whose lab part of this work was carried out. I thank Constantin Pape (Georg-August University Göttingen) for the helpful discussion; Michalis Averof for feedback on the manuscript. This research was supported by the European Research Council, under the European Union Horizon 2020 programme, grant ERC-2015-AdG #694918, and by the Agence Nationale de la Recherche of France, grant ANR-21-CE13-0044-01.

